# Cardiorespiratory fitness predicts cortical thickness of medial temporal brain areas associated with spatial cognition in young but not older adults

**DOI:** 10.1101/2021.04.11.439355

**Authors:** Michael A. Rosario, Kathryn L. Kern, Shiraz Mumtaz, Thomas W. Storer, Karin Schon

## Abstract

Cardiorespiratory fitness (CRF) has been shown to have a potent effect on neurocognitive health. However, less is known about the impact of CRF on extrahippocampal neocortical regions in the medial temporal lobes (MTL). Specifically, it is unclear whether CRF modulates these MTL regions in young adulthood and if these brain areas are differentially related to CRF in young vs. older adults. The primary goal of the current study was to investigate if CRF predicted cortical thickness of MTL neocortical regions that, along with the hippocampus, are critical for spatial learning and memory. Additionally, given the established role of the MTL cortices in spatial navigation, we sought to determine if CRF and MTL cortical thickness would predict greater subjective sense of direction in both young and older adults. Cross-sectional data from 56 young adults (20-35 years) and 44 older adults (55-85 years) were included. Using hierarchical multiple regression analyses, we confirmed significant positive relationships between greater CRF and greater left entorhinal, left parahippocampal, and left perirhinal cortical thickness in young, but not older, adults. Left parahippocampal thickness interacted with age group to differentially predict sense of direction in young and older adults. Young adults displayed a positive, and older adults a negative, relationship between left parahippocampal thickness and sense of direction. Our findings extend previous work on the association between CRF and hippocampal subfield structure in young adulthood to left MTL neocortical regions.

**Highlights:**

- Cardiorespiratory fitness assessed in young and older adults using a submaximal treadmill test.
- Surface-based structural analysis of cortical thickness of medial temporal regions.
- Cardiorespiratory fitness predicted left medial temporal cortical thickness in young but not older adults.
- Left parahippocampal thickness differentially predicted sense of direction in young and older adults.

## 1. Introduction

The medial temporal lobe (MTL) system is composed of interconnected anatomical structures, including the hippocampus (HPC), entorhinal cortex (ERC), parahippocampal cortex (PHC), and perirhinal cortex (PRC) (Squire et al., 2004). The PHC and PRC contain separate, yet slightly overlapping reciprocal connections with the ERC (Suzuki and Amaral, 1994), and the ERC has direct reciprocal connections with the HPC (Witter and Amaral, 1991). As part of a network essential for learning and memory, these MTL structures work cooperatively within a larger functional and structural extrahippocampal network (Ekstrom et al., 2017; Moffat et al., 2007) to support spatial cognition. Initial electrophysiological support for the role of the HPC in spatial learning and memory was established through the discovery of place cells (O’Keefe, 1976; O’Keefe and Dostrovsky, 1971). The MTL was further implicated in spatial navigation through the discovery of time (Eichenbaum, 2014), grid (Hafting et al., 2005), head direction (Taube et al., 1990), speed (Kropff et al., 2015), and boundary (Lever et al., 2009) cells. The PHC and PRC have been posited to underlie the representation of context and items, respectively (Squire and Zola-Morgan, 1991), with the parahippocampal place area, located within the posterior PHC, maintaining representations of spatial layout (Epstein and Kanwisher, 1998) and the PRC in more general visuospatial binding of objects within space (Connor and Knierim, 2017). Supporting this, the binding of items and contexts (BIC) model suggests that the PRC codes item information, whereas the PHC codes information for context (e.g., spatial, temporal). This item (e.g., “what”) and context (e.g. “where”) binding in PRC and PHC is then followed by a binding of the “what” and “where” of an event by the hippocampal formation (Ranganath, 2010). Considered altogether, these rodent electrophysiological studies and computational models highlight a critical role for the MTL cortices in spatial cognition.

Along with the aforementioned rodent models, non-human primate and human studies also support the role of the MTL cortices in spatial navigation. Electrophysiological recordings in monkeys showed place-related neural signals during virtual navigation in the HPC (Hori et al., 2005) and PHC (Furuya et al., 2014). Neurotoxic lesioning of the HPC in rhesus monkeys impaired spatial relational learning (Lavenex et al., 2006). Follow-up histological analyses showed that this lesioning also resulted in partial to extensive damage of the human PHC homologue in monkeys (Lavenex et al., 2006), suggesting possible PHC involvement in spatial relational learning. In treatment-resistant epileptic patients, lesions of the PHC resulted in spatial memory impairment (Ploner et al., 2000). In addition, an intracranial electrode study in epileptic patients showed that function of the left and right HPC has been shown to underlie specific components of spatial cognition, namely spatial memory for items by left HPC and spatial navigation by right HPC, in humans (Jordan, 2020; Miller et al., 2018). Functional magnetic resonance imaging (fMRI) investigations showed hippocampal recruitment during virtual navigation of a town (Maguire et al., 1998) and parahippocampal activation when participants virtually navigated a maze (Aguirre et al., 1996). A more recent fMRI study in human subjects showed macroscopic grid-cell like signals associated with the ERC during virtual navigation in an open arena (Doeller et al., 2010). Collectively, the above literature provides strong evidence for a role of the HPC and extrahippocampal MTL cortical regions in spatial cognition, including spatial navigation.

In addition to the MTLs supporting spatial navigation, these regions are exquisitely sensitive to the effects of aging on brain structure and spatial ability. In humans, older age was associated with greater HPC and PHC atrophy (Jack et al., 1997). Further, longitudinal assessments have shown age-related ERC and PHC structural atrophy in healthy adults (Daugherty and Raz, 2017; Raz et al., 2005, 2004; Shaw et al., 2016). In addition, age-related structural atrophy in these MTL regions is heterogeneous, such that the HPC atrophies at a faster rate compared to the ERC in healthy aging (Raz et al., 2005). Aging associated behavioral and functional changes in MTL-dependent spatial navigation and episodic memory also exist (Lester et al., 2017; Zhong and Moffat, 2018). Cross-sectional work has demonstrated an age-related decline in spatial mnemonic performance across the lifespan during a virtual route disambiguation task (Nauer et al., 2020b). Furthermore, two functional neuroimaging studies examining the neural correlates of spatial navigation showed reduced activation of MTL regions while navigating a virtual environment (Moffat et al., 2006) and encoding of visual cues for goal-directed spatial navigation for older compared to young adults (Antonova et al., 2009). Moreover, in the latter study, reduced bilateral PHC volume was associated with attenuated blood-oxygen-level-dependent signal of the PHC in older adults (Antonova et al., 2009). Considered together, these studies show that there are structural, behavioral, and functional differences in aging that are associated with experimental MTL-dependent measures of spatial navigation.

Whereas aging has been associated with reduced brain structural integrity and cognitive function, physical activity and aerobic exercise have been suggested as ways to mitigate these age-related changes (Hillman et al., 2008; Voss et al., 2013). In humans, aerobic exercise increases cardiorespiratory fitness (CRF) (Hagberg et al., 1989; Kohrt et al., 1991), which is a measure of one’s capacity to support ongoing physical activity through the combined efforts of the respiratory, cardiovascular, and musculoskeletal systems (ACSM, 2010). In middle-aged and older adults in particular, previous neuroimaging studies have shown profound positive associations between aerobic exercise, CRF, and neurocognitive integrity of cortical and subcortical regions (Erickson et al., 2014; Erickson and Kramer, 2009; Voelcker-Rehage and Niemann, 2013; Voss et al., 2013). In midlife, greater CRF was associated with increased bilateral PRC volume (Tian et al., 2015). Moreover, in older adults, greater CRF was associated with reduced age-related cortical atrophy across multiple brain regions, including prefrontal, inferior, and middle temporal cortices (Colcombe et al., 2003) and directly related to greater bilateral hippocampal volume (Erickson et al., 2009). Longitudinally, increase in CRF after a year-long exercise intervention (Erickson et al., 2011) was significantly associated with attenuation of age-related bilateral hippocampal volume loss in older adults. Behaviorally, using a cross-sectional design, greater CRF attenuated age-related decline across the lifespan in behavioral performance on a virtual navigation task requiring mnemonic disambiguation (Nauer et al., 2020b). Accordingly, these cross-sectional and longitudinal findings provide complementary evidence that CRF attenuates age-related decline of neocortical and subcortical neurocognitive integrity in aging.

A growing literature has shown that CRF also modulates cognitive function, MTL-dependent behavior, and brain structure in young adults. Kronman, Kern, and colleagues (2020) showed that in young adults, greater CRF predicted increased effective connectivity between the HPC and other regions of the default mode network (Kronman et al., 2020). Separately, Nauer, Dunne, et al. (2020) showed that in initially lower-fit young adults, increasing CRF through aerobic exercise training was associated with significant improvement in hippocampal-dependent memory performance (Nauer et al., 2020a). Consistent with these data, one study showed that increasing CRF through aerobic exercise training is associated with a significant improvement in visuospatial memory in young adults (Stroth et al., 2009). Beyond the demonstrated relationship between CRF and MTL-dependent function, recent work has investigated relationships between CRF and MTL structure. In a cross-sectional voxel-based morphometry study, greater CRF was associated with greater right ERC volume in young adults (Whiteman et al., 2016). Schwarb and colleagues (2017) found that greater CRF was associated with increased hippocampal integrity in young adults (Schwarb et al., 2017). In the same study, they found that hippocampal integrity mediated the relationship between CRF and performance on a spatial relational task (Schwarb et al., 2017). Furthermore, an exercise training study showed that increasing CRF through 12 weeks of aerobic exercise training resulted in increased volume of the left anterior HPC that was specific to the dentate gyrus/CA3 subfield in young adults (Nauer et al., 2020a). Complementary work by another laboratory showed that increased CRF following a six-week aerobic exercise training program was associated with increased anterior hippocampal volume in young and middle-aged adults (Thomas et al., 2016). Altogether, these findings suggest a role of CRF in influencing memory and visuospatial processes reliant on the MTL system. Furthermore, they suggest that hippocampal and entorhinal structure is positively influenced by CRF in young adulthood, rather than only in the presence of age-related atrophy, but little is known regarding other MTL neocortical structures implicated in spatial navigation.

A recent review by Voss and colleagues (2019) has suggested the use of spatial tasks as a way to more sensitively assess function of the MTL system in relation to exercise in comparison to broader assessments of cognitive function (Voss et al., 2019). A past study showed that greater CRF was associated with greater activation of a network of brain regions including the HPC, parietal cortex, and the parahippocampal gyrus during an allocentric navigation task, in middle-aged adults (Holzschneider et al., 2012), although this was not associated with increased performance on the task. Separately, measures of subjective sense of direction have been shown to be associated with greater repetition suppression within the PHC, indicating enhanced encoding of scenes over time for participants who endorsed greater sense of direction (Epstein et al., 2005). Moreover, in young adults, a structural neuroimaging study demonstrated that greater bilateral ERC and PHC volume was associated with greater self-reported sense of direction (Hao et al., 2016). Taken together, the aforementioned research raises the question of the relationship between CRF, MTL cortical thickness, and navigation ability.

The current study was designed to examine the influence of CRF upon left and right extrahippocampal MTL cortical regions implicated in spatial cognition in young and older adults. Here, we take a region of interest approach to the MTL based on the extant functional and structural literature. The primary goal of this study was to investigate the association between CRF and brain structure of MTL cortical regions that, along with the HPC, are critical for spatial cognition. Based on the above literature, we hypothesized that greater CRF should positively predict left and right ERC, PHC, and PRC thickness in young and older adulthood. Additionally, we also hypothesized that MTL cortical thickness would be positively associated with greater subjective sense of direction in young and older adults. To test our hypotheses, we used a submaximal treadmill test to estimate CRF in young and older adults and an automatic segmentation protocol to measure MTL cortical thickness. We found that greater CRF was associated with greater ERC, PHC, and PRC thickness in young, but not older, adults and that these associations were lateralized to the left hemisphere. We also showed that left PHC thickness differentially predicted sense of direction across age groups. Our findings extend a growing literature on the impact of CRF on brain structure in young and older adults from the HPC to the MTL neocortex, which are associated with spatial cognition.

## 2. Materials and Methods

### 2.1 Participants

Data for this study comes from two larger randomized controlled clinical trials (study 1: Neuroimaging Study of Exercise and Memory Function, ClinicalTrials.gov Identifier: NCT02057354) (Kern et al., 2020; Kronman et al., 2020; Nauer et al., 2020a); (study 2: The Entorhinal Cortex and Aerobic Exercise in Aging, ClinicalTrials.gov Identifier: NCT02775760) (Kern et al., 2020) which are both focused on investigating the effects of exercise on MTL structure and function. Baseline data from 56 young adults (18 - 35 years), 20 older adults (55 - 85 years) from study 1, and 24 older adults (60 - 80 years) from study 2 were included for the purposes of the current study. Participants were recruited from the greater Boston area via flyers and advertisements in local papers.

Participants underwent a prescreening process over the phone for inclusion and exclusion criteria. For young adults, exclusion criteria included severe anemia; history or occurrence of musculoskeletal, circulatory, or pulmonary conditions; diagnosis of an electrolyte disorder; acute infection, cancer, obesity as determined by American College of Sports Medicine (ACSM) guidelines (ACSM, 2010), diabetes mellitus type 1 or type 2, kidney failure, liver disease, thyroid disorders such as thyrotoxicosis/hyperthyroidism; history or diagnosis of psychiatric or neurological conditions; cardioactive or psychoactive medications; and selfreported drug abuse or alcohol misuse. Young adult women were also excluded if they were pregnant or breast-feeding. In addition to these exclusion criteria for young adults, older adults were also screened for evidence of cognitive impairment using the Dementia Rating Scale-2 (DRS-2) (Mattis, 1976). Only cognitively intact older adults were included in the study. For study 2, exclusion criteria were identical to those described for study 1 except participants with heart, circulatory, respiratory, or musculoskeletal conditions could receive clearance by their primary care physician to enter the study. Finally, both young and older adult participants were screened for MRI contraindicators (e.g., ferro-magnetic metal in or on the body that could not be removed, claustrophobia).

Inclusion criteria for both young and older adults included being generally healthy, non-smoking, and sedentary, defined as less than 30 minutes three times per week of moderate intensity physical activity over the last three months per ACSM guidelines (ACSM, 2010). Additionally, participants were fluent English speakers and had normal or corrected to normal vision. All participants signed a consent form approved by the Boston University School of Medicine Institutional Review Board and this research was conducted under the guidelines of the Declaration of Helsinki.

### 2.2 Experimental Overview

Over three study visits, participants were consented and screened, underwent cardiorespiratory fitness testing, and completed MR imaging and cognitive testing. The visits were completed within a two to three-week period, with MR imaging occurring no later than a week and not earlier than 24 hours after fitness testing to control for the acute effect of exercise on brain function and structure (Suwabe et al., 2017).

### 2.3 Assessment of Cardiorespiratory Fitness

We operationally defined CRF as aerobic capacity determined by maximal oxygen uptake 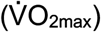. Following previous work (Kern et al., 2020; Kronman et al., 2020; Nauer et al., 2020a) we used a submaximal graded treadmill exercise test following a modified Balke protocol (ACSM, 2010) at the Boston University Fitness and Recreation Center in Boston, MA. This test protocol includes a 3-minute warm up followed by 8-12 minutes of data collection, followed by a 3-minute cool down. This protocol required participants to walk at their pre-determined fastest comfortable walking pace with an incrementally increasing grade. We monitored heart rate continuously using a heart rate monitor affixed to a chest strap (Polar, model H1) that wirelessly paired to a heart rate watch (study 1: Polar, model FT7; study 2: Polar, model A300). Using this heart rate monitor, we recorded heart rate observed in the last five seconds of every minute of testing. We measured blood pressure and recorded participants’ rating of their perceived exertion, RPE, (Borg, 1982) every 3 minutes during the exercise test in line with established guidelines (ACSM, 2010). We terminated testing when the participant reached 85% of their age-predicted maximum heart rate (Tanaka et al., 2001). Oxygen uptake 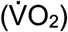 was estimated from treadmill speed and grade using standard equations which take advantage of the known linear relationship between heart rate and oxygen uptake (Wasserman, 2012) and the oxygen cost associated with walking at increasing grade, based on ACSM guidelines (Equation (1)):

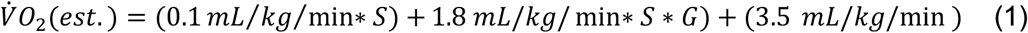

which has previously been described in detail (Kern et al., 2020; Kronman et al., 2020). In brief, 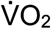 is predicted from gross oxygen uptake (mL/kg/min), treadmill speed (*S*; m/min), percent treadmill grade (*G*; treadmill grade/100), and 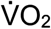 at rest (3.5mL/kg/min), respectively (ACSM, 2010). Finally, we used linear regression to predict 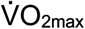 based on the known relationship between heart rate and 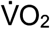 determined from work rate and the participant’s age-predicted maximum heart rate. Participants were asked not to perform any strenuous activities 24 hours prior to the test and not to consume any caffeine three hours prior to fitness testing. This exercise protocol allowed us to safely and accurately estimate 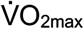 for both young and older adults (Hagberg, 1994).

### 2.4 Magnetic Resonance Image Acquisition and Image Analysis

#### 2.4.1 MRI

##### 2.4.1.1. Study 1

Participants from study 1 were scanned at the Boston University School of Medicine Center for Biomedical Imaging using a 3 T Philips Achieva scanner with an 8-channel head coil. We collected high-resolution T1-weighted structural scans (multi-planar rapidly acquired gradient echo images; SENSitivity Encoding P reduction: 1.5, S reduction: 2; TR = 6.7 ms, TE = 3.1 ms, flip angle = 9°, field of view = 25 cm, Matrix Size = 256 × 254, 150 slices, resolution = 0.98 mm × 0.98 mm × 1.22 mm).

##### 2.4.1.2. Study 2

Participants from study 2 were scanned at Boston University Cognitive Neuroimaging Center using a 3 Tesla Siemens MAGNETOM Prisma MRI scanner equipped with a stock 64-channel head coil. We collected a high-resolution whole-brain structural T1-weighted magnetization-prepared rapid acquisition gradient multi-echo (multi-echo MPRAGE volume (slices (sagittal) = 176, TR = 2200 ms, TE = 1.67 ms, TI = 1100 ms, flip angle = 7°, field of view = 230 x 230 mm, acquisition matrix = 230 x 230, voxel resolution = 1.0 mm x 1.0 mm x 1.0 mm, GRAPPA acceleration = 4).

#### 2.4.2 Regions of Interest

We conducted all automatic segmentations using FreeSurfer 6.0, a well-documented and free software available for download online (http://surfer.nmr.mgh.harvard.edu/) (Fischl, 2012). FreeSurfer is a standardized, automatic segmentation tool that constructs surface-based representations of cortical thickness calculated as the closest distance from the gray/white matter boundary to the gray/CSF boundary (Fischl and Dale, 2000). Within FreeSurfer there are no atlases that encompass all of our ROIs. Thus, measures of cortical thickness for these regions, constructed using FreeSurfer’s recon-all command, were extracted separately for the ERC (Desikan et al., 2006), PHC (Desikan et al., 2006), and PRC (Augustinack et al., 2013), for our *a priori* hypotheses detailed above. To decrease the likelihood of a Type 1 error, we restricted our analyses to a limited set of extrahippocampal cortical regions in the MTL that support spatial cognition constructed using FreeSurfer’s anatomical demarcation (Augustinack et al., 2013; Desikan et al., 2006). For each participant, we visually inspected white matter and pial surface boundaries to assure proper ROI segmentation (Dale et al., 1999). No manual correction was necessary.

### 2.5 Santa Barbara Sense of Direction Scale

We used the Santa Barbara Sense of Direction (SBSOD) scale (Hegarty et al., 2002) to investigate potential relationships between MTL cortex thickness and subjective spatial navigation. The SBSOD scale is a 15-item self-report questionnaire of one’s ability to accurately navigate an environment. Participants were asked to select how much they agreed with a statement using a 7-point Likert scale ranging from *Strongly Agree* to *Strongly Disagree*. Some example questions include: “*My “sense of direction” is very good”; “I can usually remember a new route after I have traveled it only once*”. SBSOD scale scores were calculated as the averaged sum of all items after reverse-scoring positively phrased items. Higher scores indicate better subjective spatial ability. Three participants (1 young adult and 2 older adults from Study 1) had missing SBSOD questionnaire data and were excluded from these analyses.

### 2.6 Statistical Analyses

Statistical analyses were conducted using R (4.0.0) and RStudio (1.2.5042). We tested our primary outcome variables for normality using the Shapiro-Wilk test. All were normally distributed in the overall sample and within the young and older adult samples separately. Demographic characteristics were calculated using independent sample’s t-test. Continuous variables were summarized by mean, range, and standard deviation. Sex was summarized by percentage.

First, we conducted young and older adult group comparisons using an analysis of covariance (ANCOVA) method controlling for sex and education to establish if there was a differential impact of age group on ROI thickness. Next, we used multiple regression analyses that included a CRF (operationalized as estimated 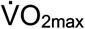) by age group interaction term to determine whether CRF predicted left or right ERC, PHC, and PRC thickness differentially between young and older adults, holding sex and education constant. Subsequently, we used ordinary least squares multiple regression models to test our primary hypothesis that greater CRF would positively predict greater left or right MTL ROI thickness in our young and older adults, separately, holding sex, chronological age, and education constant. In our analyses investigating sense of direction, we included an age group by ROI interaction term to determine whether cortical thickness of our ROI predicted SBSOD scores differentially between age groups. We then separated our analyses by age group to investigate SBSOD scores from our primary ROIs in separate models, holding sex, chronological age, and education constant, and included a ROI x estimated 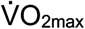 interaction term to determine if CRF had any appreciable impact on the relationship between ROI and sense of direction. A significant interaction effect signified that estimated 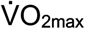 moderated the ROI slope on sense of direction beyond the solitary influence of either estimated 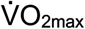 or ROI thickness, in the statistical model.

Continuous predictor variables were standardized by mean-centering and scaling by 2 standard deviations(Gelman, 2008). ΔR^2^, calculated using the *getDeltaRsquare* function in R, was used as a measure of effect size to determine the variance explained by the interaction effect and the inclusion of estimated 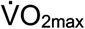 in our models. We corrected for multiple comparisons with the False Discovery Rate (FDR) method in R using the *p-adjust* function, and statistical significance of p-values was considered at the corrected level of *p_FDR_* < .05.

## 3. Results

### 3.1 Participant Characteristics

Participant characteristics are described in Table 1. DRS-2 raw memory and total scores are provided to show that our older adult sample is cognitively healthy compared with normative data(Johnson-Greene, 2004), and are not included in further analyses. We additionally provided a visual representation of the distribution of estimated 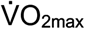 and SBSOD scores by age group and sex (see Figure 1).

**Table 1.** Participant Characteristics. Parametric data are presented as Mean (SD). Categorical data are presented using percentages. Differences between age groups statistically significant at *p* < .05 * or *p* < .001 ***. YA: Young adults; OA: Older Adults; 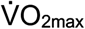 (ml/kg/min): maximal oxygen uptake; SBSOD: Santa Barbara Sense of Direction Scale; DRS-2: Dementia Rating Scale 2.

| Demographics | Range<sub>YA</sub> | Mean<sub>YA</sub> | Range<sub>OA</sub> | Mean<sub>OA</sub> |
| --- | --- | --- | --- | --- |
| N = 100 |  | N = 56 |  | N = 44 |
|  |  | M(SD) |  | M(SD) |
| Age (years) | 20 - 35 | 25.9 (3.5) | 55 - 85 | 65.6 (7.3)* |
| Sex | - | 71.4% female | - | 57.8% female |
| Education (years) | 12 - 20 | 16.8 (2.6) | 12 - 20 | 17 (2.2) |
| Estimated $\dot{V}O_{2max}$ (ml/kg/min) | 25.1 - 57.5 | 36.7 (6.8) | 17.7 - 42.7 | 30.4 (5.9)* |
| Estimated $\dot{V}O_{2max}$ Percentile | 3 - 90 | 41.7 (22.64) | 27 - 99 | 78.3 (21.13) |
| SBSOD | 1.7 - 6.3 | 4.4(1.1) | 2.8 - 67 | 5.1 (.87)*** |
| DRS-2 Total Raw Score | - | - | 133 - 144 | 140.97 (2.26) |
| DRS-2 Memory Raw Score | - | - | 22 - 25 | 24.18 (.99) |

**Figure 1.**
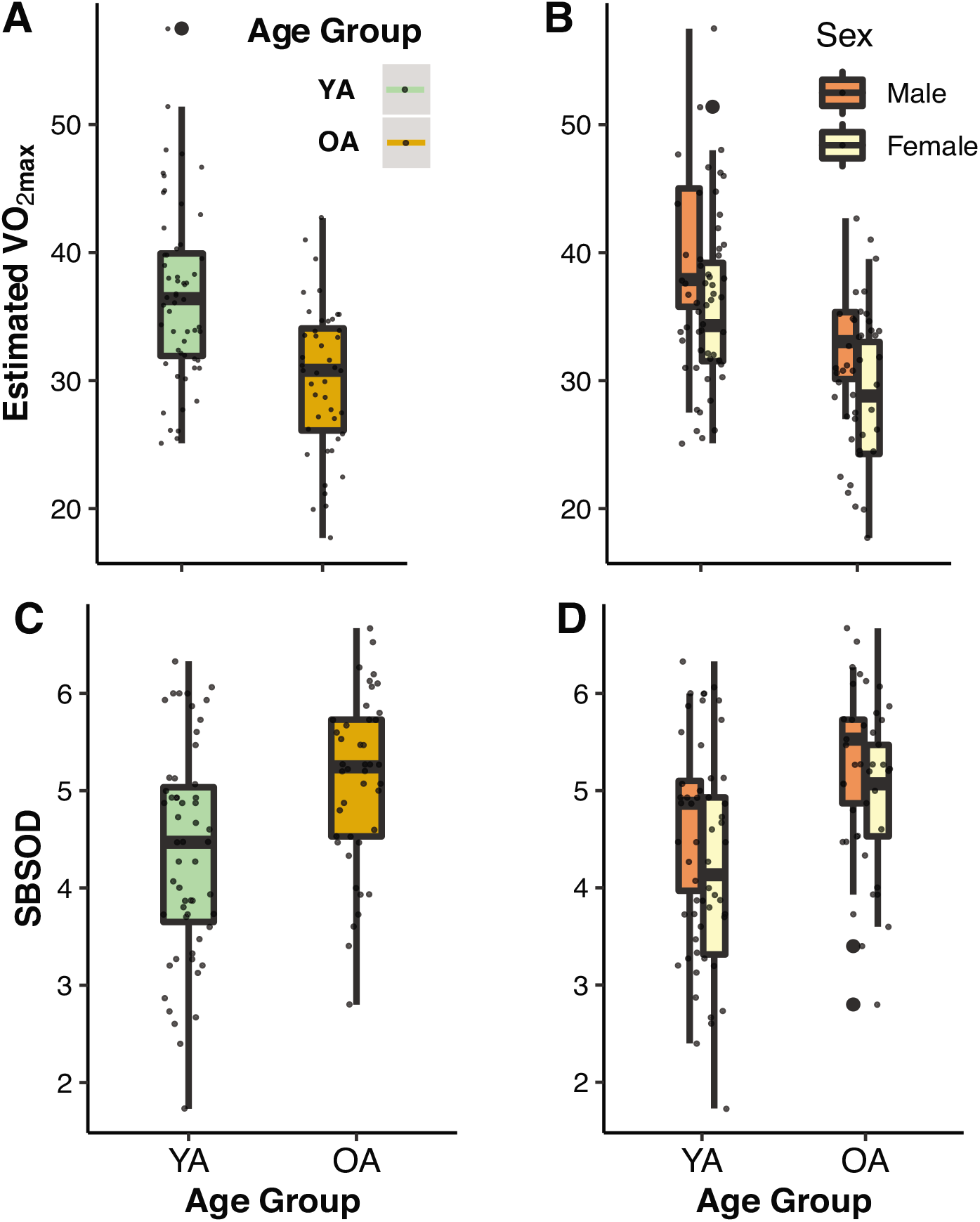
Boxplots of distributions by age group and sex for CRF and SBSOD. Distribution of estimated 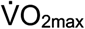 by A) age group and by B) age group and sex. Distribution of subjective sense of direction scores by C) age group and by D) age group and sex. SBSOD: Santa Barbara Sense of Direction, YA: Young Adult, OA: Older Adult: Estimated 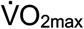: cardiorespiratory fitness operationalized. (See article online for color version of figure).

### 3.2 Association between cardiorespiratory fitness and cortical thickness differs between young and older adults

To understand our findings within the context of the existing CRF, aging, and brain structure literature, we first assessed whether there were mean differences in MTL cortical thickness, independent of CRF, by age group. Using an ANCOVA controlling for sex and education, we compared young and older adults and found that there was no effect of age group on the thickness of the left ERC (*F*(3,97) = .981, *p* = .33), right ERC (*F*(3,97) = 1.19, *p* = .28), or left PRC (*F*(3,97) = 2.35, *p* = .13). However, there was a significant effect of age group on thickness of the left PHC (*F*(3,97) = 10.25, *p* < .01), right PHC (*F*(3,97) = 9.10, *p* < .01), and right PRC (*F*(3,97) = 4.29, *p* = .04). The nature of these significant differences was such that young adults demonstrated significantly greater mean cortical thickness than older adults (see Figure 2).

**Figure 2.**
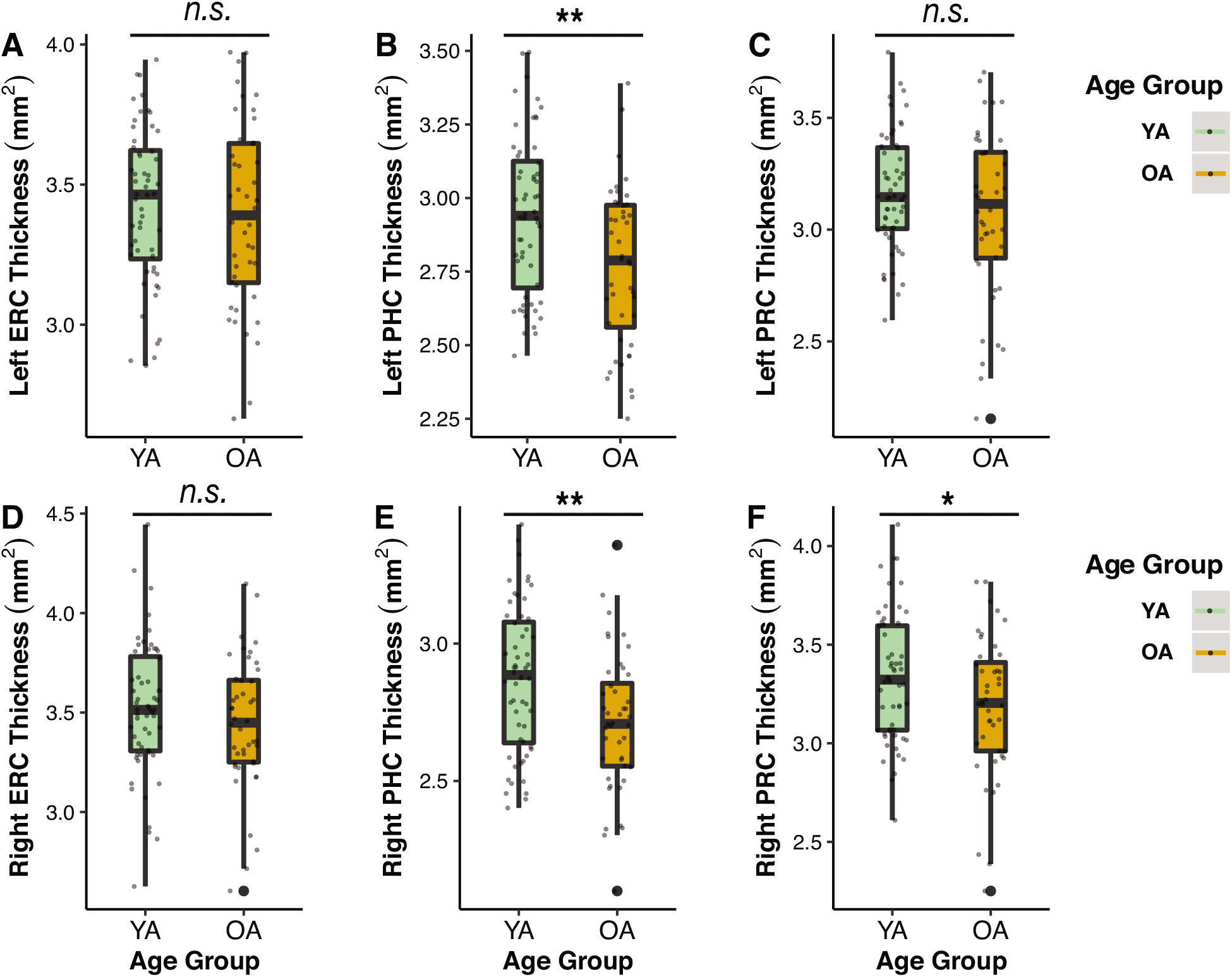
Box plots displaying significant mean differences between young and older adults by ROI. Top row presents left hemisphere ROIs: A) ERC, B) PHC, C) PRC; bottom row presents right hemisphere ROIs: D) ERC, E) PHC, and F) PRC, by age group, independent of the effect of cardiorespiratory fitness. ERC: entorhinal cortex, PHC: parahippocampal cortex; PRC: perirhinal cortex; YA: Young Adult; OA: Older Adult. * *p* < .05, ** *p* < .01. (See article online for color version of figure).

Subsequently, using ordinary least squares regression, controlling for sex and education, we tested for a CRF x Age Group interaction effect on left and right ERC, PHC, and PRC thickness. Although we predicted similar relationships for young and older adults, we conducted these analyses in order to determine if CRF would modulate extrahippocampal MTL cortical thickness differently in young and older adults as seen in previous work (Williams et al., 2017). We found no significant CRF x Age Group interaction effects on the thickness of the left ERC (*ß* = -.25 CI[-.53,.03], *t*(95) = −1.80, *p_FDR_* = .11, Δ*R^2^* = .01; model: *F*(5,94) = 1.94, *R^2^* = .09;), right ERC (*ß* = -.12 CI[-.44,.20], *t*(95) = -.73, *p_FDR_* = .47, Δ*R^2^* < .01; model: *F*(5,94) = .70, *R^2^* = .04), left PHC (*ß* = -.32, CI[-.57,-.07], *t*(95) = −2.57, *p_FDR_* = .07, Δ*R^2^* = .06; model: *F*(5,94) = 4.07, *R^2^* = .18), right PHC (*ß* = -.27, CI[-.51,-.03], *t*(95) = −2.20, *p_FDR_* = .09, Δ*R^2^* = .04; model: *F*(5,94) = 3.29, *R^2^* = .15), left PRC (*ß* = -.29 CI[-.56,-.01], *t*(95) = −2.04, *p_FDR_* = .09, Δ*R^2^* = .04; model: *F*(5,94) = 4.21, *R^2^* = .18), or right PRC (*ß* = -.19 CI[-.51,.13], *t*(95) = −1.19, *p_FDR_* = .28, Δ*R^2^* = .01; model: *F*(5,94) = 1.67, *R^2^* = .08) after correction for multiple comparisons.

We next investigated whether CRF predicted left or right ERC, PHC, or PRC thickness separately in young or older adults, first without the inclusion of CRF (model 1) and then with the inclusion of CRF (model 2). In order to do so, we separated our dataset by age group and controlled for chronological age, sex, and education within our models. In contrast to our hypothesis that greater CRF would predict increased cortical integrity in older adults, there were no significant relationships between greater CRF and left nor right ERC, PHC, or PRC thickness for older adults (see Table 2 for statistics). However, in agreement with our hypothesis in our young adult sample, greater CRF was positively correlated with greater left ERC, left PHC, and left PRC thickness (see Table 2; see Figure 3). There was no significant association between CRF and right ROI in young adults. CRF explained an additional 17%, 9%, and 11% of the variance for left ERC, left PHC, and left PRC thickness, respectively, beyond chronological age, sex, and education, in these statistical models. These analyses confirmed a positive association between greater CRF and increased MTL ROI thickness within the young adult group, and this was specific to the left hemisphere.

**Table 2.**
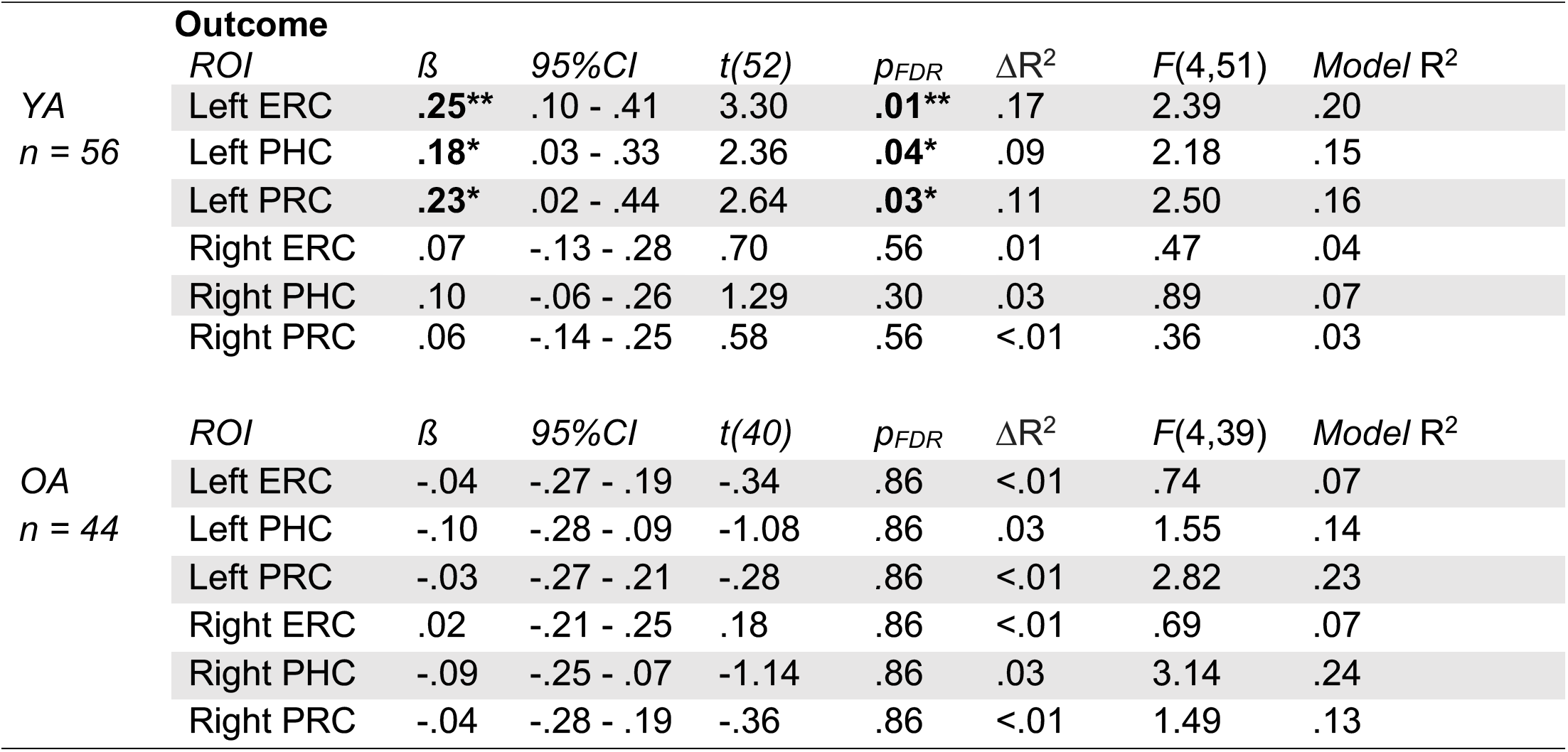
Hierarchical multiple linear regression results for CRF and cortical thickness by age group. Table 2 details the change in variance explained by CRF (estimated 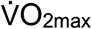) in separate models for young and older adults by ROI, controlling for chronological age, education, and sex. Significant results are bolded. OA: Older Adults; YA: Young Adults; ROI: region of interest, CI: confidence interval, ERC: entorhinal cortex, PHC: parahippocampal cortex, PRC: perirhinal cortex. * *p* < .05, ** *p* < .01

**Figure 3.**
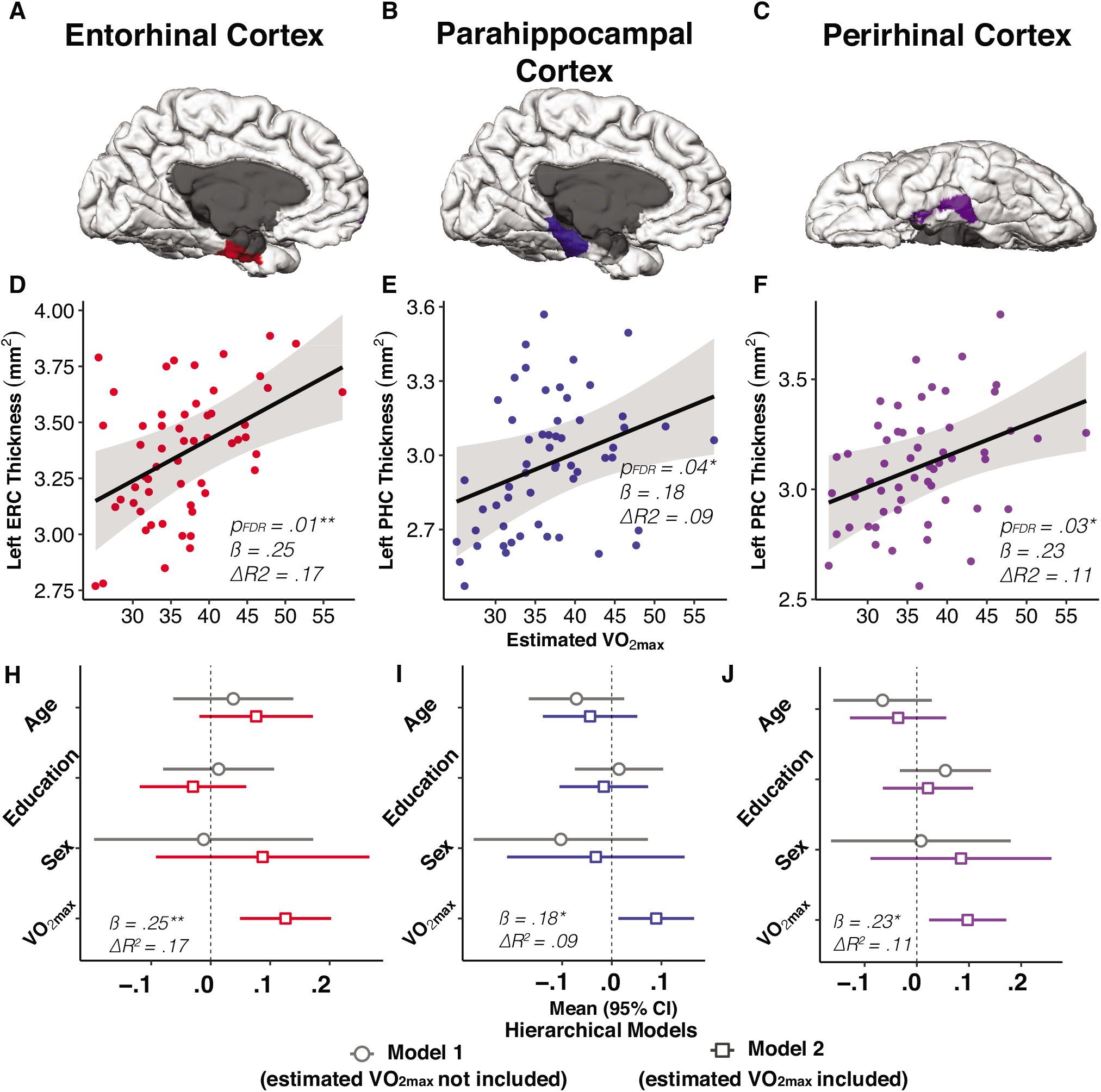
Greater CRF is associated with left ROI thickness in young adults. Figures within the top row present an inflated pial, medial surface view of the region of interest from a representative participant for each corresponding scatterplot: A) entorhinal cortex (red), B) parahippocampal cortex (blue), C) perirhinal cortex (purple). Partial residual plots displaying the regression results for left D) ERC, E) PHC, and F) PRC thickness are presented in the second row, respectively, controlling for chronological age, education, and sex. For each corresponding scatterplot, we used a forest plot to display the overall results of the hierarchical linear model for each ROI, including covariates. Model 1, without the inclusion of estimated 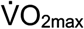, is represented in with an open circle, and model 2, with the inclusion of estimated 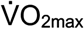, is represented by an open square. The *p_FDR_*, beta coefficient, and Δ*R^2^* are presented for each significant model. (See article online for color version of figure).

### 3.3 Left parahippocampal thickness differentially predicts poorer sense of direction for young and older adults

Based on our above finding that there was a significant difference in SBSOD scores by age group whereby older adults self-rated greater sense of direction compared to young adults (*t*(95) = −3.64, *p* < .001), we first investigated whether age group interacted with left and right ERC, PHC, and PRC thickness or CRF to predict sense of direction. Thus, we included a CRF x Age Group and a ROI x Age Group interaction term in separate models, controlling for sex and education. There was no significant CRF x Age Group interaction effect on SBSOD scores (*ß* = -.17, CI[-1.15,.82], *t*(97) = -.34, *p_FDR_* = .78), thus we excluded CRF from all further analyses.

There was a significant Left PHC thickness x Age Group interaction effect on SBSOD scores (*ß* = −1.28, CI[-2.14,-.42], *t*(97) = −2.94, *p_FDR_* = .03, Δ*R*^2^ = .08; model: *F*(5,91) = 4.80, *R*^2^ = .21) (see Figure 4A). For all other regions there were no significant ROI x Age Group interactions. We focused our subsequent analyses on the left PHC in order to determine whether the significant Left PHC thickness x Age Group interaction was driven by young and/or older adults, using multiple regression analyses within separate age groups. We found a significant positive relationship between greater left PHC thickness and greater SBSOD scores (*ß* = .59, CI[.001,1.18], *t*(52) = 2.01, *p* = .05, Δ*R*^2^ = .07; model: *F*(3,51) = 2.0, *R*^2^ = .11), controlling for sex and education, in young adults (see Figure 4B). Contrary to our prediction, in older adults there was a significant negative relationship between greater left PHC thickness and lower SBSOD scores (*ß* = -.63, CI[-1.19,-.06], *t*(39) = −2.24, *p* = .03, Δ*R*^2^ = .11; model: *F*(3,38) = 1.88, *R*^2^ = .13), controlling for sex and education (see Figure 4C). Hence, our Left PHC thickness x Age Group interaction effect shows that greater sense of direction was associated with lower left PHC thickness in older adults and greater left PHC thickness in young adults.

**Figure 4.**
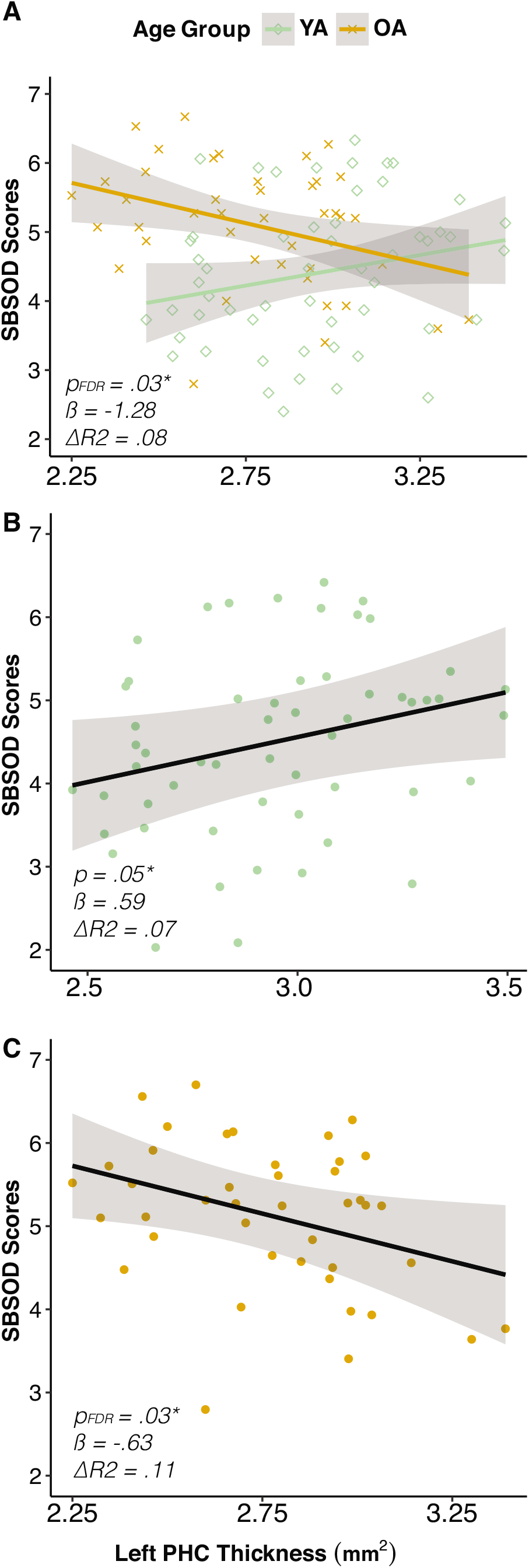
Santa Barbara Sense of Direction Scale Scores and Left Parahippocampal Cortical Thickness. A) Interaction model: There was a significant Left PHC Thickness x Age Group interaction effect on subjective sense of direction. B) Subjective sense of direction was positively associated with greater left PHC thickness, in young adults, controlling for sex and education. C) In older adults, subjective sense of direction was negatively associated with left PHC thickness, controlling for sex and education. YA: Young Adult; OA: Older Adult; PHC: Parahippocampal Cortex. SBSOD: Santa Barbara Sense of Direction. (See article online for color version of figure).

## 4.0 Discussion

The goal of the current study was to investigate the relationship between CRF and structural integrity of MTL regions that subserve spatial cognition using a surface-based structural analysis of cortical thickness. We tested the hypothesis that CRF would positively predict left and right ERC, PHC, and PRC thickness in young and older adults. First, we investigated whether there was a significant difference in MTL neocortical thickness in young compared to older adults. We found that there was a significant difference in left PHC, right PHC, and right PRC thickness between young and older adults, such that young adults had greater cortical thickness in these regions. In line with our primary hypothesis for our young adult sample, at higher CRF, ERC, PHC, and PRC thickness was greater, but unexpectedly, were lateralized to the left hemisphere. Contrary to our hypothesis for our older adult sample, CRF did not predict MTL ROI cortical thickness. We then investigated whether a subjective assessment of sense of direction was related to CRF and to cortical thickness in our ROIs. We found that age group and left PHC thickness significantly interacted to predict sense of direction. Ensuing analyses within age group, specific to the left PHC, showed that young adults displayed a positive relationship and older adults a negative relationship between left PHC thickness and sense of direction. These findings extend a growing literature on the association between CRF and brain structure in young adults, by focusing on structural measures in extrahippocampal regions. Specifically, we show that these regions, of the left hemisphere, are sensitive to CRF-associated cortical plasticity in young, but not older, adults, and provide data suggesting that CRF may modulate extrahippocampal regions of the MTL in the absence of age-related structural decline.

### 4.1 Cardiorespiratory fitness predicts neocortical thickness in the MTL in young but not older adults

The primary objective of this study was to investigate the modulating role of CRF on extrahippocampal neocortical regions of the MTL implicated in spatial cognition in both young and older adults. Although we hypothesized a positive association between CRF and MTL ROI in both young and older adults, we first conducted analyses investigating whether age and CRF would interact to differentially predict left or right ERC, PHC, and PRC thickness similarly to previous work (Williams et al., 2017). There were no significant interactions between aerobic capacity and age group on left or right ERC, PHC, or PRC thickness. We then tested our primary *a priori* hypotheses by stratifying our following analyses by young and older adults to determine if CRF predicted cortical thickness of MTL ROIs within each age group.

Contrary to our hypothesis, there were no significant relationships between aerobic capacity and MTL neocortical thickness in our older adult sample. Considering the previously published literature, recent studies in older adults have investigated the effects of exercise on brain structural integrity and have largely focused on hippocampal volume. This body of work has demonstrated that greater CRF (Erickson et al., 2009), greater physical activity (Erickson et al., 2010), and aerobic exercise training that increases CRF (Erickson et al., 2011) are associated with increased hippocampal volume and/or attenuation of age-related hippocampal atrophy. In a separate study conducted in older adults, increased CRF was associated with increased hippocampal perfusion and hippocampal head volume after undergoing a three-month exercise intervention (Maass et al., 2015). A recently published study focused on the relationship between CRF and hippocampal subfield volume in healthy older adults, showed that CRF was significantly related to bilateral hippocampal subiculum volume, and that this effect was driven by the women in this sample (Kern et al., 2020). Although we recruited participants who identified as sedentary (ACSM, 2010), the participants in our older adult sample were generally more fit in comparison to national fitness norms (Myers et al., 2017). This could potentially explain this lack of relationship, given that previous findings showed an increase in left anterior dentate gyrus volume following twelve weeks of exercise training was dependent on being initially lower fit (Nauer et al., 2020a). Although this study was done in young adults, these results suggests that the biggest benefit of CRF may be conferred to lower-fit individuals who have the capacity to increase fitness levels. Additionally, the above literature gives further indication to the lack of association identified in our findings, potentially signifying either a sex-specific or hippocampal subfield-specific impact of CRF on brain structure in aging.

In contrast to our findings in older adults, our results in young adults complement a growing body of literature on the positive relationships between CRF and MTL structure in young adults (Nauer et al., 2020a; Schwarb et al., 2017; Whiteman et al., 2016). Our analyses showed that greater CRF had a moderate to large effect on left ERC, left PHC, and left PRC thickness in young adults. Previously, using a voxel intensity-based morphometry method, greater CRF was shown to predict greater right ERC volume in a sample of young adults (Whiteman et al., 2016). Using a surface-based analysis approach, we extended these findings from ERC volume to left ERC thickness in young adults. Additionally, we extended our understanding of the relationship between CRF and MTL ROI thickness to the left PHC and left PRC. In conjunction with the aging literature described above, these data in young adulthood suggest that the ERC, and other regions of the MTL including the PHC and PRC, may be associated with CRF in the absence of age-related structural atrophy. Thus, our data builds on and complements a growing literature on the relationship between CRF and MTL structure beyond the HPC to the left ERC, left PHC, and left PRC in young adulthood.

Because we had no *a priori* hypotheses regarding laterality, we did not conduct analyses directly comparing hemispheres. However, it is of note that the relationships observed in young adults between CRF and MTL cortical thickness were found in the left, not right, hemisphere. In studies of physical activity in older adults, where CRF is not measured, longitudinal and prospective research has shown a specific effect of physical activity on left lateralized brain structures, specifically in the left HPC (Erickson et al., 2010; Rosano et al., 2017). A meta-analysis of randomized controlled trials of CRF-induced changes following exercise training across the adult lifespan also reported a specific effect of aerobic exercise on left, but not right, HPC (Firth et al., 2018). Longitudinal work in young adults showed that a 12-week exercise intervention resulted in a volume increase in dentate gyrus/CA3 in the hippocampal head that was specific to the left hemisphere (Nauer et al., 2020a). In contrast to these studies, an exploratory cross-sectional study investigating the relationship between CRF and cortical thickness in healthy young vs. older adults found a more global impact of CRF in both the left and right hemispheres (Williams et al., 2017). Our current findings on the relationships between CRF and left ERC, left PHC, and left PRC thickness add to our growing understanding of the impact of exercise and fitness on brain structure, and in the context of the extant literature, suggest that there may be a preferential impact on the left hemisphere.

Our understanding of the underlying neurobiological mechanisms modulated by CRF in humans is still limited. Research in rodents provide suitable theoretical grounding on the impact of CRF on brain structure and function, and have demonstrated that growth and genetic factors may underlie the current study’s observed associations between CRF and MTL cortices. For example, in rodents, the HPC and ERC showed greater expression of growth factors after wheel running which, in turn, were involved in the survival and differentiation of existing neurons (Ickes et al., 2000; Pham et al., 1999). In addition, independent of exercise effects, rodents from strains that engaged in more voluntary wheel running showed higher levels of brain-derived neurotrophic factor (BDNF), a neurochemical growth factor important for neuronal survival and synaptic plasticity, in the MTL system, at rest (Johnson et al., 2003; Johnson and Mitchell, 2003). This finding indicates a potential genetic predisposition to exercise effects on brain structure and function. Complementary to this research, a recent human neuroimaging study in male adolescents investigated whether individual differences in BDNF genotype would moderate the impact of CRF on brain structures. They found such a moderation effect whereby participants with a BDNF genotype that facilitates greater BDNF expression showed a significant positive relationship between CRF and bilateral medial precuneus surface area (Herting et al., 2016). This suggests the need for future studies to control for genetic factors in order to distinguish between exercise effects and genetic effects of CRF.

In addition to the research in rodents suggesting that growth and genetic factors may be modulated by CRF, work focused on morphological changes in both rodents and humans have proposed that neurogenesis, myelination, and vascularization may similarly be targets of increased CRF. Early research on wheel-running in rodents showed a preferential impact of exercise on neurogenesis in the dentate gyrus subfield of the HPC (Van Praag et al., 1999b, 1999a). More recent work using diffusion tensor imaging has shown an exercise-induced change in hippocampal myelination in both humans and rodents (Islam et al., 2020; Thomas et al., 2016). Separately, in a study focusing on the impact of exercise training on the MTLs, Pereira and colleagues (2007) found increased blood flow to the dentate gyrus subregion of the HPC in young mice after two weeks of voluntary wheel running (Pereira et al., 2007). In the same study, they found a corresponding increase in cerebral blood flow to the dentate gyrus in a small sample of young to middle-aged adults, as well as a non-significant positive impact on increased cerebral blood flow to the ERC, after a three-month exercise intervention (Pereira et al., 2007), although this non-significant effect may be due in part to their small sample size. Altogether, these neurobiological mechanisms provide different targets that may be modulated by CRF and necessitate additional multimodal and concurrent cross-species investigations to determine whether the observed changes associated with CRF are indeed conserved across species.

### 4.2 Left parahippocampal cortex is differentially associated with sense of direction in young and older adults

We next investigated the relationship between CRF and our ROIs with sense of direction. We found that there was a significant interaction between age group and left PHC thickness predicting subjective sense of direction. The PHC has been implicated in both allocentric and egocentric navigation (Colombo et al., 2017; Epstein, 2008; Epstein et al., 2017), and is also related to subjective sense of direction (Hao et al., 2016). Thus, we asked whether left PHC thickness predicted sense of direction in our sample and found that in young adults, greater left PHC thickness predicted better sense of direction, whereas in older adults, greater left PHC thickness predicted poorer sense of direction. These results in our young adult sample extend previous work, which showed that greater bilateral PHC volume was positively correlated with higher SBSOD scores in a large sample of young adults (Hao et al., 2016). Moreover, in young adults, Antonova and colleagues (2009) found that there was greater left parahippocampal gyrus functional activity during encoding of an allocentric spatial navigation task, although this was not related to behavioral outcomes (Antonova et al., 2009). In contrast to our findings in young adults, our results in older adults are contradictory to previous research in spatial navigation and aging. A previous study in older adults showed that reduced parahippocampal gyrus volume predicted reduced signal in this region during encoding of virtual cues in an allocentric spatial navigation task (Antonova et al., 2009), suggesting that structural integrity of the PHC may underlie successful navigational ability in older adults. We found no significant interaction between CRF and age group on sense of direction in our study. One study in young adults that did investigate the influence of CRF on spatial performance found that a sensitive measure of hippocampal integrity, its viscoelasticity, mediated the relationship between CRF and a spatial relational task (Schwarb et al., 2017). This suggests that CRF may be more sensitively related to objective measurements of spatial navigation. Further neuroimaging studies that examine both brain structure and function are needed to determine whether CRF modulates or mediates brain function and the underlying neural correlates that support spatial navigation ability in young and older adults. Further empirical research is also needed to investigate whether CRF may rescue functional and behavioral impairment in navigation ability in older, relative to young, adults.

### Limitations

A major strength of the study is the relatively large sample size compared to similar work investigating relationships between aerobic capacity and structural integrity between young and older adults. One limitation that exists is the use of submaximal instead of maximal testing to estimate 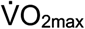. We chose this protocol to ensure the safety of our older participants during exercise testing, and used the same protocol for our young adults to maintain concordance between age groups. Using submaximal tests has been established as a safe way to reliably estimate 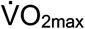, compared to high-intensity maximal tests, to ensure safety of populations with different mobility needs (ACSM, 2010; Hagberg, 1994). Our older adults were also higher fit compared to national norms (Myers et al., 2017). Future studies should examine whether the benefit of increasing aerobic capacity on brain structure and function is inversely proportional to the CRF level of the individual at the beginning of an exercise training program.

Although we cannot determine causality within our cross-sectional analyses, we showed that CRF was positively associated with left hemispheric MTL neocortical thickness in young adults. Randomized controlled trials that measure the change in extrahippocampal cortical thickness across multiple time points would be able to examine the nature of causal influence among our correlational findings. Our current study lacked any objective measures of virtual navigation performance to supplement our measure of subjective spatial ability. Future studies should collect experimental data of spatial navigation to understand how brain-CRF relationships support performance on spatial navigation tasks, and whether CRF mitigates age-related and neuropathological decline in spatial cognition (Maass and Shine, 2019). A methodological limitation of the work includes the potential spatial overlap between our measures of ERC, PHC, and PRC thickness, given they were constructed from two separate anatomical atlases within the MTL (Augustinack et al., 2013; Desikan et al., 2006). However, previous work has shown that automatic segmentation protocols ensure increased replicability and reliability (Han et al., 2006) by moving beyond the bias introduced during manual segmentation.

### Conclusion

The current study extends the extant literature by providing additional evidence for the modulatory role of CRF on cortical thickness in young adults. More specifically, these data extend previous work on the relationship between CRF and volume of the ERC in young adults, to left ERC, left PHC, and left PRC thickness, suggesting that these regions may be amenable to experience-dependent structural plasticity outside of age-related structural atrophy. Additionally, although there is a rich literature supporting the positive effect of exercise and CRF in older adults, our results observed in young adults did not extend to our older adult sample. Our findings on subjective sense of direction showed that left PHC was positively associated with sense of direction in young, but negatively associated with sense of direction in older adults. These paradoxical results in older adults require further investigation with objective measures of spatial navigation ability. Across the lifespan there are potentially different neurobiological mechanisms that CRF may influence at different timepoints in the adult lifespan, thus longitudinal research may provide the experimental insight needed to understand the neurobiological underpinnings of CRF and exercise effects on brain structure and function across the lifespan.

## Acknowledgements

This work was supported in part by the National Institutes of Health R00 AG036845 (KS), R21 AG049968 (KS), a Robert Wood Johnson Foundation Health Policy Research Scholars grant (MAR), and the Boston University Clinical and Translational Science Institute (UL-TR000157). We thank Rachel Nauer Wehr and Matthew Dunne for data collection. We also thank additional members of the Brain Plasticity and Neuroimaging Laboratory involved in data collection and Lucas Carstensen at the Center for Systems Neuroscience for his feedback on the manuscript.

## Contributions

K.S. and M.A.R. designed the study and developed the methodology. M.A.R. analyzed the data and wrote the first draft of the manuscript. K.L.K. and S.M. collected the data for Study 2. K.L.K, S.M, T.S., & K.S. provided critical revision of the manuscript.

## Data Availability Statement

Data available upon reasonable request and upon establishment of a formal data sharing agreement.

## References

ACSM, 2010. ACSM’s Guidelines for exercise Testing and Prescripstion. Prim. Care 21, 55–67. https://doi.org/10.1249/MSS.0b013e318213fefb

Aguirre, G.K., Detre, J.A., Alsop, D.C., D’Esposito, M., 1996. The parahippocampus subserves topographical learning in man. Cereb. Cortex 6, 823–829. https://doi.org/10.1093/cercor/6.6.823

Antonova, E., Parslow, D., Brammer, M., Dawson, G.R., Jackson, S.H.D., Morris, R.G., 2009. Age-related neural activity during allocentric spatial memory. Memory 17, 125–143. https://doi.org/10.1080/09658210802077348

Augustinack, J.C., Huber, K.E., Stevens, A.A., Roy, M., Frosch, M.P., van der Kouwe, A.J.W., Wald, L.L., Van Leemput, K., McKee, A.C., Fischl, B., 2013. Predicting the location of human perirhinal cortex, Brodmann’s area 35, from MRI. Neuroimage 64, 32–42. https://doi.org/10.1016/j.neuroimage.2012.08.071

Borg, G.A.V., 1982. Psychophysical bases of perceived exertion. Med. Sci. Sports Exerc. 14, 377–381. https://doi.org/10.1249/00005768-198205000-00012

Colcombe, S.J., Erickson, K.I., Raz, N., Webb, A.G., Cohen, N.J., McAuley, E., Kramer, A.F., 2003. Aerobic fitness reduces brain tissue loss in aging humans. Journals Gerontol. - Ser. A Biol. Sci. Med. Sci. 58, 176–180. https://doi.org/10.1093/gerona/58.2.m176

Colombo, D., Serino, S., Tuena, C., Pedroli, E., Dakanalis, A., Cipresso, P., Riva, G., 2017. Egocentric and allocentric spatial reference frames in aging: A systematic review. Neurosci. Biobehav. Rev. https://doi.org/10.1016/j.neubiorev.2017.07.012

Connor, C.E., Knierim, J.J., 2017. Integration of objects and space in perception and memory. Nat. Neurosci. https://doi.org/10.1038/nn.4657

Dale, A.M., Fischl, B., Sereno, M.I., 1999. Cortical surface-based analysis: I. Segmentation and surface reconstruction. Neuroimage 9, 179–194. https://doi.org/10.1006/nimg.1998.0395

Daugherty, A.M., Raz, N., 2017. A virtual water maze revisited: Two-year changes in navigation performance and their neural correlates in healthy adults. Neuroimage 146, 492–506. https://doi.org/10.1016/j.neuroimage.2016.09.044

Desikan, R.S., Ségonne, F., Fischl, B., Quinn, B.T., Dickerson, B.C., Blacker, D., Buckner, R.L., Dale, A.M., Maguire, R.P., Hyman, B.T., Albert, M.S., Killiany, R.J., 2006. An automated labeling system for subdividing the human cerebral cortex on MRI scans into gyral based regions of interest. Neuroimage 31, 968–980. https://doi.org/10.1016/j.neuroimage.2006.01.021

Doeller, C.F., Barry, C., Burgess, N., 2010. Evidence for grid cells in a human memory network. Nature 463, 657–661. https://doi.org/10.1038/nature08704

Eichenbaum, H., 2014. Time cells in the hippocampus: A new dimension for mapping memories. Nat. Rev. Neurosci. https://doi.org/10.1038/nrn3827

Ekstrom, A.D., Huffman, D.J., Starrett, M., 2017. Interacting networks of brain regions underlie human spatial navigation: A review and novel synthesis of the literature. J. Neurophysiol. https://doi.org/10.1152/jn.00531.2017

Epstein, R., Kanwisher, N., 1998. A cortical representation the local visual environment. Nature. https://doi.org/10.1038/33402

Epstein, R.A., 2008. Parahippocampal and retrosplenial contributions to human spatial navigation. Trends Cogn. Sci. https://doi.org/10.1016/j.tics.2008.07.004

Epstein, R.A., Higgins, J.S., Thompson-Schill, S.L., 2005. Learning places from views: Variation in scene processing as a function of experience and navigational ability. J. Cogn. Neurosci. https://doi.org/10.1162/0898929052879987

Epstein, R.A., Patai, E.Z., Julian, J.B., Spiers, H.J., 2017. The cognitive map in humans: Spatial navigation and beyond. Nat. Neurosci. https://doi.org/10.1038/nn.4656

Erickson, K.I., Kramer, A.F., 2009. Aerobic exercise effects on cognitive and neural plasticity in older adults. Br. J. Sports Med. https://doi.org/10.1136/bjsm.2008.052498

Erickson, K.I., Leckie, R.L., Weinstein, A.M., 2014. Physical activity, fitness, and gray matter volume. Neurobiol. Aging. https://doi.org/10.1016/j.neurobiolaging.2014.03.034

Erickson, K.I., Prakash, R.S., Voss, M.W., Chaddock, L., Hu, L., Morris, K.S., White, S.M., Wójcicki, T.R., McAuley, E., Kramer, A.F., 2009. Aerobic fitness is associated with hippocampal volume in elderly humans. Hippocampus 19, 1030–1039. https://doi.org/10.1002/hipo.20547

Erickson, K.I., Raji, C.A., Lopez, O.L., Becker, J.T., Rosano, C., Newman, A.B., Gach, H.M., Thompson, P.M., Ho, A.J., Kuller, L.H., 2010. Physical activity predicts gray matter volume in late adulthood: The Cardiovascular Health Study. Neurology 75, 1415–1422. https://doi.org/10.1212/WNL.0b013e3181f88359

Erickson, K.I., Voss, M.W., Prakash, R.S., Basak, C., Szabo, A., Chaddock, L., Kim, J.S., Heo, S., Alves, H., White, S.M., Wojcicki, T.R., Mailey, E., Vieira, V.J., Martin, S.A., Pence, B.D., Woods, J.A., McAuley, E., Kramer, A.F., 2011. Exercise training increases size of hippocampus and improves memory. Proc. Natl. Acad. Sci. U. S. A. 108, 3017–3022. https://doi.org/10.1073/pnas.1015950108

Firth, J., Stubbs, B., Vancampfort, D., Schuch, F., Lagopoulos, J., Rosenbaum, S., Ward, P.B., 2018. Effect of aerobic exercise on hippocampal volume in humans: A systematic review and meta-analysis. Neuroimage. https://doi.org/10.1016/j.neuroimage.2017.11.007

Fischl, B., 2012. FreeSurfer. Neuroimage. https://doi.org/10.1016/j.neuroimage.2012.01.021

Fischl, B., Dale, A.M., 2000. Measuring the thickness of the human cerebral cortex from magnetic resonance images. Proc. Natl. Acad. Sci. U. S. A. 97, 11050–11055. https://doi.org/10.1073/pnas.200033797

Furuya, Y., Matsumoto, J., Hori, E., Boas, C.V., Tran, A.H., Shimada, Y., Ono, T., Nishijo, H., 2014. Place-related neuronal activity in the monkey parahippocampal gyrus and hippocampal formation during virtual navigation. Hippocampus 24, 113–130. https://doi.org/10.1002/hipo.22209

Gelman, A., 2008. Scaling regression inputs by dividing by two standard deviations. Stat. Med. https://doi.org/10.1002/sim.3107

Hafting, T., Fyhn, M., Molden, S., Moser, M.B., Moser, E.I., 2005. Microstructure of a spatial map in the entorhinal cortex. Nature 436, 801–806. https://doi.org/10.1038/nature03721

Hagberg, J.M., 1994. Exercise assessment of arthritic and elderly individuals. Baillieres. Clin. Rheumatol. 8, 29–52. https://doi.org/10.1016/S0950-3579(05)80223-7

Hagberg, J.M., Graves, J.E., Limacher, M., Woods, D.R., Leggett, S.H., Cononie, C., Gruber, J.J., Pollock, M.L., 1989. Cardiovascular reponses of 70-to 79-yr-old men and women to exercise training. J. Appl. Physiol. 66, 2589–2594. https://doi.org/10.1152/jappl.1989.66.6.2589

Han, X., Jovicich, J., Salat, D., van der Kouwe, A., Quinn, B., Czanner, S., Busa, E., Pacheco, J., Albert, M., Killiany, R., Maguire, P., Rosas, D., Makris, N., Dale, A., Dickerson, B., Fischl, B., 2006. Reliability of MRI-derived measurements of human cerebral cortical thickness: The effects of field strength, scanner upgrade and manufacturer. Neuroimage. https://doi.org/10.1016/j.neuroimage.2006.02.051

Hao, X., Huang, Y., Li, X., Song, Y., Kong, X., Wang, X., Yang, Z., Zhen, Z., Liu, J., 2016. Structural and functional neural correlates of spatial navigation: a combined voxel-based morphometry and functional connectivity study. Brain Behav. 6, e00572. https://doi.org/10.1002/brb3.572

Hegarty, M., Richardson, A.E., Montello, D.R., Lovelace, K., Subbiah, I., 2002. Development of a self-report measure of environmental spatial ability. Intelligence 30, 425–447. https://doi.org/10.1016/S0160-2896(02)00116-2

Herting, M.M., Keenan, M.F., Nagel, B.J., 2016. Aerobic fitness linked to cortical brain development in adolescent males: Preliminary findings suggest a possible role of BDNF genotype. Front. Hum. Neurosci. 10, 327. https://doi.org/10.3389/fnhum.2016.00327

Hillman, C.H., Erickson, K.I., Kramer, A.F., 2008. Be smart, exercise your heart: Exercise effects on brain and cognition. Nat. Rev. Neurosci. https://doi.org/10.1038/nrn2298

Holzschneider, K., Wolbers, T., Röder, B., Hötting, K., 2012. Cardiovascular fitness modulates brain activation associated with spatial learning. Neuroimage 59, 3003–3014. https://doi.org/10.1016/j.neuroimage.2011.10.021

Hori, E., Nishio, Y., Kazui, K., Umeno, K., Tabuchi, E., Sasaki, K., Endo, S., Ono, T., Nishijo, H., 2005. Place-related neural responses in the monkey hippocampal formation in a virtual space. Hippocampus 15, 991–996. https://doi.org/10.1002/hipo.20108

Ickes, B.R., Pham, T.M., Sanders, L.A., Albeck, D.S., Mohammed, A.H., Granholm, A.C., 2000. Long-term environmental enrichment leads to regional increases in neurotrophin levels in rat brain. Exp. Neurol. 164, 45–52. https://doi.org/10.1006/exnr.2000.7415

Islam, M.R., Luo, R., Valaris, S., Haley, E.B., Takase, H., Chen, Y.I., Dickerson, B.C., Schon, K., Arai, K., Nguyen, C.T., Wrann, C.D., 2020. Diffusion tensor-MRI detects exercise-induced neuroplasticity in the hippocampal microstructure in mice. Brain Plast. https://doi.org/10.3233/bpl-190090

Jack, C.R., Petersen, R.C., Xu, Y.C., Waring, S.C., O’Brien, P.C., Tangalos, E.G., Smith, G.E., Ivnik, R.J., Kokmen, E., 1997. Medial temporal atrophy on MRI in normal aging and very mild Alzheimer’s disease. Neurology 49, 786–794. https://doi.org/10.1212/WNL.49.3.786

Johnson-Greene, D., 2004. Dementia Rating Scale-2 (DRS-2) By P.J. Jurica, C.L. Leitten, and S. Mattis: Psychological Assessment Resources, 2001. Arch. Clin. Neuropsychol. 19, 145–147. https://doi.org/10.1016/j.acn.2003.07.003

Johnson, R.A., Mitchell, G.S., 2003. Exercise-induced changes in hippocampal brain-derived neurotrophic factor and neurotrophin-3: Effects of rat strain. Brain Res. https://doi.org/10.1016/S0006-8993(03)03039-7

Johnson, R.A., Rhodes, J.S., Jeffrey, S.L., Garland, T., Mitchell, G.S., 2003. Hippocampal brain-derived neurotrophic factor but not neurotrophin-3 increases more in mice selected for increased voluntary wheel running. Neuroscience. https://doi.org/10.1016/S0306-4522(03)00422-6

Jordan, J.T., 2020. The rodent hippocampus as a bilateral structure: A review of hemispheric lateralization. Hippocampus. https://doi.org/10.1002/hipo.23188

Kern, K.L., Storer, T.W., Schon, K., 2020. Cardiorespiratory fitness, hippocampal subfield volumes, and mnemonic discrimination task performance in aging. Hum. Brain Mapp. https://doi.org/10.1002/hbm.25259

Kohrt, W.M., Malley, M.T., Coggan, A.R., Spina, R.J., Ogawa, T., Ehsani, A.A., Bourey, R.E., Martin, W.H., Holloszy, J.O., 1991. Effects of gender, age, and fitness level on response of VO2max to training in 60-71 yr olds. J. Appl. Physiol. 71, 2004–2011. https://doi.org/10.1152/jappl.1991.71.5.2004

Kronman, C.A., Kern, K.L., Nauer, R.K., Dunne, M.F., Storer, T.W., Schon, K., 2020. Cardiorespiratory fitness predicts effective connectivity between the hippocampus and default mode network nodes in young adults. Hippocampus 30, 526–541. https://doi.org/10.1002/hipo.23169

Kropff, E., Carmichael, J.E., Moser, M.B., Moser, E.I., 2015. Speed cells in the medial entorhinal cortex. Nature 523, 419–424. https://doi.org/10.1038/nature14622

Lavenex, P.B., Amaral, D.G., Lavenex, P., 2006. Hippocampal lesion prevents spatial relational learning in adult macaque monkeys. J. Neurosci. 26, 4546–4558. https://doi.org/10.1523/JNEUROSCI.5412-05.2006

Lester, A.W., Moffat, S.D., Wiener, J.M., Barnes, C.A., Wolbers, T., 2017. The Aging Navigational System. Neuron. https://doi.org/10.1016/j.neuron.2017.06.037

Lever, C., Burton, S., Jeewajee, A., O’Keefe, J., Burgess, N., 2009. Boundary vector cells in the subiculum of the hippocampal formation. J. Neurosci. 29, 9771–9777. https://doi.org/10.1523/JNEUROSCI.1319-09.2009

Maass, A., Düzel, S., Goerke, M., Becke, A., Sobieray, U., Neumann, K., Lövden, M., Lindenberger, U., Bäckman, L., Braun-Dullaeus, R., Ahrens, D., Heinze, H.J., Müller, N.G., Düzel, E., 2015. Vascular hippocampal plasticity after aerobic exercise in older adults. Mol. Psychiatry. https://doi.org/10.1038/mp.2014.114

Maass, A., Shine, J.P., 2019. Navigating the future of clinical assessments. Brain. https://doi.org/10.1093/brain/awz121

Maguire, E.A., Burgess, N., Donnett, J.G., Frackowiak, R.S.J., Frith, C.D., O’Keefe, J., 1998. Knowing where and getting there: A human navigation network. Science (80-.). 280, 921–924. https://doi.org/10.1126/science.280.5365.921

Mattis, S., 1976. Mental status examination for organic mental syndrome in the elderly patient, in: Geriatric Psychiatry.

Miller, J., Watrous, A.J., Tsitsiklis, M., Lee, S.A., Sheth, S.A., Schevon, C.A., Smith, E.H., Sperling, M.R., Sharan, A., Asadi-Pooya, A.A., Worrell, G.A., Meisenhelter, S., Inman, C.S., Davis, K.A., Lega, B., Wanda, P.A., Das, S.R., Stein, J.M., Gorniak, R., Jacobs, J., 2018. Lateralized hippocampal oscillations underlie distinct aspects of human spatial memory and navigation. Nat. Commun. https://doi.org/10.1038/s41467-018-04847-9

Moffat, S.D., Elkins, W., Resnick, S.M., 2006. Age differences in the neural systems supporting human allocentric spatial navigation. Neurobiol. Aging 27, 965–972. https://doi.org/10.1016/j.neurobiolaging.2005.05.011

Moffat, S.D., Kennedy, K.M., Rodrigue, K.M., Raz, N., 2007. Extrahippocampal contributions to age differences in human spatial navigation. Cereb. Cortex 17, 1274–1282. https://doi.org/10.1093/cercor/bhl036

Myers, J., Kaminsky, L.A., Lima, R., Christle, J.W., Ashley, E., Arena, R., 2017. A Reference Equation for Normal Standards for VO2 Max: Analysis from the Fitness Registry and the Importance of Exercise National Database (FRIEND Registry). Prog. Cardiovasc. Dis. https://doi.org/10.1016/j.pcad.2017.03.002

Nauer, R.K., Dunne, M.F., Stern, C.E., Storer, T.W., Schon, K., 2020a. Improving fitness increases dentate gyrus/CA3 volume in the hippocampal head and enhances memory in young adults. Hippocampus 30, 488–504. https://doi.org/10.1002/hipo.23166

Nauer, R.K., Schon, K., Stern, C.E., 2020b. Cardiorespiratory fitness and mnemonic discrimination across the adult lifespan. Learn. Mem. 27, 91–103. https://doi.org/10.1101/lm.049197.118

O’Keefe, J., 1976. Place units in the hippocampus of the freely moving rat. Exp. Neurol. https://doi.org/10.1016/0014-4886(76)90055-8

O’Keefe, J., Dostrovsky, J., 1971. The hippocampus as a spatial map. Brain Res.

Pereira, A.C., Huddleston, D.E., Brickman, A.M., Sosunov, A.A., Hen, R., McKhann, G.M., Sloan, R., Gage, F.H., Brown, T.R., Small, S.A., 2007. An in vivo correlate of exercise-induced neurogenesis in the adult dentate gyrus, Proceedings of the National Academy of Sciences of the United States of America. PNAS. https://doi.org/10.1073/pnas.0611721104

Pham, T.M., Ickes, B., Albeck, D., Söderström, S., Granholm, A.C., Mohammed, A.H., 1999. Changes in brain nerve growth factor levels and nerve growth factor receptors in rats exposed to environmental enrichment for one year. Neuroscience 94, 279–286. https://doi.org/10.1016/S0306-4522(99)00316-4

Ploner, C.J., Gaymard, B.M., Rivaud-Péchoux, S., Baulac, M., Clémenceau, S., Samson, S., Pierrot-Deseilligny, C., 2000. Lesions affecting the parahippocampal cortex yield spatial memory deficits in humans. Cereb. Cortex 10, 1211–1216. https://doi.org/10.1093/cercor/10.12.1211

Ranganath, C., 2010. Binding items and contexts: The cognitive neuroscience of episodic memory. Curr. Dir. Psychol. Sci. https://doi.org/10.1177/0963721410368805

Raz, N., Lindenberger, U., Rodrigue, K.M., Kennedy, K.M., Head, D., Williamson, A., Dahle, C., Gerstorf, D., Acker, J.D., 2005. Regional brain changes in aging healthy adults: General trends, individual differences and modifiers. Cereb. Cortex 15, 1676–1689. https://doi.org/10.1093/cercor/bhi044

Raz, N., Rodrigue, K.M., Head, D., Kennedy, K.M., Acker, J.D., 2004. Differential aging of the medial temporal lobe: A study of a five-year change. Neurology 62, 433–438. https://doi.org/10.1212/01.WNL.0000106466.09835.46

Rosano, C., Guralnik, J., Pahor, M., Glynn, N.W., Newman, A.B., Ibrahim, T.S., Erickson, K., Cohen, R., Shaaban, C.E., MacCloud, R.L., Aizenstein, H.J., 2017. Hippocampal Response to a 24-Month Physical Activity Intervention in Sedentary Older Adults. Am. J. Geriatr. Psychiatry. https://doi.org/10.1016/j.jagp.2016.11.007

Schwarb, H., Johnson, C.L., Daugherty, A.M., Hillman, C.H., Kramer, A.F., Cohen, N.J., Barbey, A.K., 2017. Aerobic fitness, hippocampal viscoelasticity, and relational memory performance. Neuroimage. https://doi.org/10.1016/j.neuroimage.2017.03.061

Shaw, M.E., Sachdev, P.S., Anstey, K.J., Cherbuin, N., 2016. Age-related cortical thinning in cognitively healthy individuals in their 60s: The PATH Through Life study. Neurobiol. Aging 39, 202–209. https://doi.org/10.1016/j.neurobiolaging.2015.12.009

Squire, L.R., Stark, C.E.L., Clark, R.E., 2004. The medial temporal lobe. Annu. Rev. Neurosci. 27, 279–306. https://doi.org/10.1146/annurev.neuro.27.070203.144130

Squire, L.R., Zola-Morgan, S., 1991. The medial temporal lobe memory system. Science (80-.). https://doi.org/10.1126/science.1896849

Stroth, S., Hille, K., Spitzer, M., Reinhardt, R., 2009. Aerobic endurance exercise benefits memory and affect in young adults. Neuropsychol. Rehabil. 19, 223–243. https://doi.org/10.1080/09602010802091183

Suwabe, K., Hyodo, K., Byun, K., Ochi, G., Fukuie, T., Shimizu, T., Kato, M., Yassa, M.A., Soya, H., 2017. Aerobic fitness associates with mnemonic discrimination as a mediator of physical activity effects: Evidence for memory flexibility in young adults. Sci. Rep. 7, 1–10. https://doi.org/10.1038/s41598-017-04850-y

Suzuki, W.A., Amaral, D.G., 1994. Topographic organization of the reciprocal connections between the monkey entorhinal cortex and the perirhinal and parahippocampal cortices. J. Neurosci. https://doi.org/10.1523/jneurosci.14-03-01856.1994

Tanaka, H., Monahan, K.D., Seals, D.R., 2001. Age-predicted maximal heart rate revisited. J. Am. Coll. Cardiol. 37, 153–156. https://doi.org/10.1016/S0735-1097(00)01054-8

Taube, J.S., Muller, R.U., Ranck, J.B., 1990. Head-direction cells recorded from the postsubiculum in freely moving rats. I. Description and quantitative analysis. J. Neurosci. 10, 420–435. https://doi.org/10.1523/jneurosci.10-02-00420.1990

Thomas, A.G., Dennis, A., Rawlings, N.B., Stagg, C.J., Matthews, L., Morris, M., Kolind, S.H., Foxley, S., Jenkinson, M., Nichols, T.E., Dawes, H., Bandettini, P.A., Johansen-Berg, H., 2016. Multi-modal characterization of rapid anterior hippocampal volume increase associated with aerobic exercise. Neuroimage 131, 162–170. https://doi.org/10.1016/j.neuroimage.2015.10.090

Tian, Q., Studenski, S.A., Resnick, S.M., Davatzikos, C., Ferrucci, L., 2015. Midlife and Late-Life Cardiorespiratory Fitness and Brain Volume Changes in Late Adulthood: Results from the Baltimore Longitudinal Study of Aging. Journals Gerontol. - Ser. A Biol. Sci. Med. Sci. 71, 124–130. https://doi.org/10.1093/gerona/glv041

Van Praag, H., Christie, B.R., Sejnowski, T.J., Gage, F.H., 1999a. Running enhances neurogenesis, learning, and long-term potentiation in mice. Proc. Natl. Acad. Sci. U. S. A. 96, 13427–13431. https://doi.org/10.1073/pnas.96.23.13427

Van Praag, H., Kempermann, G., Gage, F.H., 1999b. Running increases cell proliferation and neurogenesis in the adult mouse dentate gyrus. Nat. Neurosci. 2, 266–270. https://doi.org/10.1038/6368

Voelcker-Rehage, C., Niemann, C., 2013. Structural and functional brain changes related to different types of physical activity across the life span. Neurosci. Biobehav. Rev. https://doi.org/10.1016/j.neubiorev.2013.01.028

Voss, M.W., Soto, C., Yoo, S., Sodoma, M., Vivar, C., van Praag, H., 2019. Exercise and Hippocampal Memory Systems. Trends Cogn. Sci. https://doi.org/10.1016/j.tics.2019.01.006

Voss, M.W., Vivar, C., Kramer, A.F., van Praag, H., 2013. Bridging animal and human models of exercise-induced brain plasticity. Trends Cogn. Sci. https://doi.org/10.1016/j.tics.2013.08.001

Wasserman, K., 2012. Principles of exercise testing and interpretation: Including pathophysiology and clinical applications, 5th ed. Wolters Kluwer Health/Lippincott Williams & Wilkins, Philadelphia.

Whiteman, A.S., Young, D.E., Budson, A.E., Stern, C.E., Schon, K., 2016. Entorhinal volume, aerobic fitness, and recognition memory in healthy young adults: A voxel-based morphometry study. Neuroimage 126, 229–238. https://doi.org/10.1016/j.neuroimage.2015.11.049

Williams, V.J., Hayes, J.P., Forman, D.E., Salat, D.H., Sperling, R.A., Verfaellie, M., Hayes, S.M., 2017. Cardiorespiratory fitness is differentially associated with cortical thickness in young and older adults. Neuroimage 146, 1084–1092. https://doi.org/10.1016/j.neuroimage.2016.10.033

Witter, M.P., Amaral, D.G., 1991. Entorhinal cortex of the monkey: V. Projections to the dentate gyrus, hippocampus, and subicular complex. J. Comp. Neurol. https://doi.org/10.1002/cne.903070308

Zhong, J.Y., Moffat, S.D., 2018. Extrahippocampal contributions to age-related changes in spatial navigation ability. Front. Hum. Neurosci. 12, 272. https://doi.org/10.3389/fnhum.2018.00272

